# Tissue Usage Preference and Intrinsically Disordered Region Remodeling of Alternative Splicing Derived Proteoforms in the Heart

**DOI:** 10.1101/2023.10.08.561375

**Authors:** Boomathi Pandi, Stella Brenman, Alexander Black, Dominic C. M. Ng, Edward Lau, Maggie P. Y. Lam

**Affiliations:** Department of Medicine/Division of Cardiology, University of Colorado School of Medicine Aurora, CO 80045, USA; Department of Biochemistry & Molecular Genetics, University of Colorado School of Medicine Aurora, CO 80045, USA; Consortium for Fibrosis Research and Translation (CFReT), University of Colorado School of Medicine Aurora, CO 80045, USA

## Abstract

A computational analysis of mass spectrometry data was performed to uncover alternative splicing derived protein variants across chambers of the human heart. Evidence for 216 non-canonical isoforms was apparent in the atrium and the ventricle, including 52 isoforms not documented on SwissProt and recovered using an RNA sequencing derived database. Among non-canonical isoforms, 29 show signs of regulation based on statistically significant preferences in tissue usage, including a ventricular enriched protein isoform of tensin-1 (TNS1) and an atrium-enriched PDZ and LIM Domain 3 (PDLIM3) isoform 2 (PDLIM3-2/ALP-H). Examined variant regions that differ between alternative and canonical isoforms are highly enriched in intrinsically disordered regions, and over two-thirds of such regions are predicted to function in protein binding and/or RNA binding. The analysis here lends further credence to the notion that alternative splicing diversifies the proteome by rewiring intrinsically disordered regions, which are increasingly recognized to play important roles in the generation of biological function from protein sequences.

## Introduction

Alternative splicing contributes to the functional diversity of the genome by allowing multiple transcript variants to be encoded by one gene. Prior analysis of alternative splicing in transcriptomics data has suggested that alternative transcripts are highly tissue specific and have strong potential to alter protein interaction networks (1–3). In contrast, proteome-level investigations of the function and regulation of alternative splicing have been lagging, presenting a significant knowledge gap in the field. Whether the majority of alternative splicing transcripts exist as stable proteins remains unclear, which has led to questions on whether alternative splicing may diversify protein function (4, 5), or instead act to modulate protein levels (6, 7). Not every transcript isoform carries equal protein coding potential as many contain frameshifts and premature stop codons that can trigger gene product degradation following transcription and/or translation; hence identifying the splice variant at the protein level is an important part to adjudicating the molecular function of the splice products. The identification of alternative proteins is at present uncommonly performed due to both analytical and computational challenges, including the possible aforementioned frameshift, low abundance of alternative transcripts, and sequence properties near exon-exon junctions (8–11). In part because of these reasons, a common practice for bottom-up proteomics studies is to report only one protein per gene or collapsing peptides across isoforms into the same protein group.

Publicly available mass spectrometry data presents opportunities to mine, analyze, and interpret existing data sets using new computational advances to derive new insights not apparent from the original study and analysis. We and others have previously employed a proteogenomics approach that combines information from transcriptomics and proteomics to identify hundreds of alternative isoforms in mammalian tissues (9, 12, 13) and cell lines (11, 14). Toward the raw mass spectrometry data, we applied a computational pipeline we developed to create custom protein sequence databases from ENCODE RNA sequencing data. We showed that such approaches can recapture non-canonical protein isoforms that are recorded by mass spectrometry but not identified in the original published studies.

The human heart is among the organs that are most heavily regulated by alternative splicing derived isoforms (9, 15, 16). Alternative splicing is an important form of gene regulation in the heart and disruption of splicing can causally lead to non-compaction and other congenital cardiomyopathies (17, 18). The expression of multiple cardiac genes is known to show biased expression across atrial and ventricular tissues, including classically known atrial and ventricular isoforms of myosin light chains 1 (MLC-1a/v) and 2 (MLC-2a/v), atrial natriuretic peptide (NPPA), and others within the heart (19). Both bottom-up (20, 21) and top-down (22, 23) mass spectrometry-based proteomics have been utilized to discover chamber specific or chamber biased proteins, which may help to explain the metabolic and functional differences between the atrium and the ventricle and shed light on the pathology of chamber specific cardiac diseases.

Here we performed a quantitative re-analysis of chamber biased expression of cardiac proteins using two deep mass spectrometry data sets that distinguished atrial and ventricular samples. We found that alternative splice variant proteoforms with chamber-enriched expression can be found in the human heart across atrial and ventricular samples. Moreover, by applying newly available deep learning strategies, we showed that alternative protein isoforms broadly rewire IDRs in the proteome and alter regions with predicted propensity for protein binding and RNA binding. The analysis here adds to the emerging view of the close connections between alternative splicing, intrinsically disordered regions (IDR), and post-translational modifications (PTMs) with protein function and contributes further evidence to the importance of alternative splicing to generating proteome diversity.

## Methods

### Mass spectrometry data re-analysis

The mass spectrometry data of human atrial and ventricular samples from 7 donors were originally described in Linscheid et al, and were acquired on a Q-Exactive HF Orbitrap instrument (Thermo) using data-dependent acquisition, with 120,000 MS1 and 30,000 MS2 resolution. The raw data were retrieved from ProteomeXchange (24) (PXD008722) using the rpx package, then converted to mzml peak lists using ThermoRawFileParser v.1.2.0 (25). To perform database search, we downloaded the UniProt SwissProt human database (2023_02 release) appended with contaminant proteins and decoy sequences (20,455 forward sequences) using the default settings in Philosopher v.4.8.1 (26). The data were then searched using MSFragger v.3.8 (27) with the default closed search settings with the following parameters: precursor_mass_lower –20 ppm, precursor_mass_upper 20 ppm, precursor_true_tolerance 20 ppm, fragment_mass_tolerance 20 ppm, isotope_error 0/1/2, num_enzyme_termini 1. The search results were post-processed using PeptideProphet, InterProphet, and ProteinProphet, followed by protein inference using the Picked Protein algorithm in Philosopher v.4.8.1 (26). Label-free quantity was calculated with match-between-runs using IonQuant v.1.9.8 (28) with the following arguments: --mbrrttol5 --mbrtoprun 10 –mbrmincorr 0.15 --ionfdr 0.1. The results are filtered for proteins to consider only proteins with multiple testing adjusted protein probability above 95% for quantification. To verify isoform identification, the mzml files were further searched using identical databases as above using Comet (v2022_01) (29) with typical settings for FT/FT spectra (peptide mass tolerance: 10 ppm; isotope error: 0/1/2/3, fragment bin tol: 0.02), followed by post-processing and FDR calculation using Percolator (Crux 4.1 distribution) (30) with default settings. Peptide spectrum matches with posterior error probability < 0.01 were considered significant.

### Custom protein isoform database generation

Isoform-aware search was performed against a UniProt SwissProt human canonical + isoform sequence database (2023_02 release) appended with contaminant proteins and decoy sequences (42,352 forward sequences). The databases were further appended with custom translated isoform sequences derived from RNA-seq data. Briefly, RNA-seq reads were retrieved from ENCODE (31) human heart total RNA-seq experiments and aligned to GRCh38 using STAR v.2.7.6a (32). Splice junctions were collated using rMATS v.4.1.0 (33), the then filtered by read counts and translated in silico using JCAST v.0.3.4 (34), requiring 16 total splice junction read counts to filter out low-read splice junctions (35).

### Differential expression analysis

The summed label-free intensity of each protein in each sample in the filtered IonQuant output was normalized using variance stabilizing transformation (36). Left atrium and right atrium samples were labeled as atrial samples and left ventricle samples were labeled as ventricular samples as in the original publications. Samples from the 7 donors were encoded as in the original publication. Differential protein abundance across atrial and ventricular samples were compared using limma v.3.52.4 (37) in R v.4.2.1 with a ∼0 + Chamber + Individual formula to contrast across chambers while adjusting for individual donors. A Benjamini-Hochberg adjusted limma p value of ≤ 0.05 for the canonical search and ≤ 0.1 for the canonical + isoform search was considered significant.

### Procedure for structure and functional prediction

Functional prediction of PDLIM3 isoforms was carried out on bases of structural and sequence using various computational methods. Protein domains and interaction partners were retrieved using InterPro (38) and StringDB v12 (39). The sequence input of each PDLIM3 isoform was input to the UCSF ChimeraX platform (40) for getting the AlphaFold2 proposed structure (41). The IDR regions were examined using Metapredict v2 (42) and their functional site is predicted by flDPnn docker version (accessed 2023-08-01) (43). The protein interaction site for AlphaFold2 proposed model was examined using ScanNet web server (accessed 2023-08-15) (44). Further, protein-protein interactions between PDLIM3 and ACTN2 were explored using AlphaFold-Multimer (45).

## Results

### Discovery of non-canonical protein isoforms with a proteogenomics pipeline

We generated a custom database for human heart protein and protein isoforms using a splice junction centric translation tool JCAST we previously developed (9). Using a mixture model to filter out low read junctions, JCAST translated a total of 2,388 non-canonical isoforms in-frame as the “tier 1” output (i.e., translated using the Ensembl annotated reading frame without encountering frameshifts or premature termination codons), from which we removed 361 titin isoforms, as their inclusion caused the ProteinProphet step of the database search to fail. Of the remaining 2,027 non-canonical sequences, 1,309 were not contained within Uniprot SwissProt canonical + isoform sequences. This custom database was merged with the Philosopher-annotated SwissProt canonical + isoform database to harmonize naming convention and yield a final database with 43,661 forward sequences. Some of the sequences are contained in TrEMBL. Thus this method allows appending of variant sequences based on likelihood of existence in a sample while limiting the inflation of database size that can come with the use of a very permissive database (9) (e.g., TrEMBL, RefSeq, or 3-frame in silico translation).

We asked whether the alternative protein isoforms may show evidence of functional regulation by considering chamber differences in expression. To do so, we used MSFragger/IonQuant label free quantification of MS1 intensity, followed by limma to compare ventricular and atrial expression of isoforms in Linscheid et al. (PXD008722) (21). The original study by Linscheid et al. found 741 proteins with significant differences in abundance between the atrial and ventricular tissues of 7 human hearts. The publication used MaxQuant and Perseus for protein identification and quantification on a SwissProt with isoform database (accessed 2016-03-07 and containing 42,138 total entries), but did not distinguish any isoform difference in their atrial vs. ventricular protein abundance comparisons.

We first sought to confirm that we could recapitulate the original study using a label free quantification workflow against a UniProt SwissProt canonical protein sequence database (2023_02 release), which contains only one canonical sequence per gene and does not contain isoform sequences. The re-analysis quantified 4,073 proteins across all samples, which recapitulated the clear separation between ventricular and atrial samples in the first principal components of protein intensity. We found 814 proteins to be significantly differentially expressed after multiple testing correction (5% FDR; 1,137 proteins at 10% FDR), including known atrial and ventricular enriched proteins and those reported in the study by Linscheid et al. These include MLC-2a (MYL7) which in the re-analysis is 99.2-fold higher in the atrium than the ventricle (adjusted p: 5.4e-11) and MLC-2v (MYL2) which is 6.4 fold higher in the ventricle (adjusted p: 9.4e–4). Similarly, other highlighted atrial-enriched proteins (MYH4, MYL4, TMOD2) and ventricular-enriched proteins (MYH7, MYL3, DSP) were significantly differentially expressed in the re-analysis. Overall, the re-analysis found a similar number of differentially expressed proteins as in the original publication (814 in this study vs. 741) at identical significance cutoff (5% FDR), despite using a different protein identification and quantification workflow (MSFragger + IonQuant vs. MaxQuant + Perseus) and sequence database (UniProt 2023_02 canonical vs. UniProt 2016 canonical + isoforms). (**Table S1: Canonical Search)**

We next searched for protein isoform expression by performing a second database search. We searched the mass spectrometry data using a custom protein database constructed from appending UniProt SwissProt canonical + isoform sequences with additional RNA-seq derived protein sequences using a proteogenomics pipeline we previously developed (9) (see Methods). We did not observe significant false discovery rate inflation from database size. Instead, the inclusion of additional isoform sequences in the database led to similar numbers of quantifiable proteins (4,284 isoforms from 4,030 genes compared between ventricles and atria with 99% probability (unique + razor peptides); compared to 4,073 in the canonical search).

The new search retained the clear ventricular and atrial separation in PC1 (**Figure 1A**). We found 760 (unique + razor peptides) proteins to be differentially expressed at 5% FDR (1,076 at 10% FDR) (**Figure 1B**) including 83 genes where more than one protein isoforms from a gene were quantified and where at least one shows a significant difference in ventricular vs. atrial distributions at 10% FDR. When only unique peptides are quantified, 29 non-canonical isoforms show significant chamber-biased expression (**Table 1; Diff. Isoforms; Table S2 full isoform search table**). We note that the requirement for unique peptides in isoform analysis is expected to lead to substantial data dropout and can also lead to some unwarranted complications: because of the piecewise nature of bottom-up proteomics similar to short-read RNA-seq, the definition of uniqueness depends on the sequence database being used. Therefore, depending on the coverage and precision of the in-silico database, a peptide may be considered non-unique because they are shared by two overlapping isoforms that are not canonical; some of the isoforms may not in fact exist as full-length translated molecules in the sample, whereas proteins with complex splicing patterns will tend to contain few unique peptides from trypsin digestion. The quantified protein ratios will also differ because fewer peptides are now admissible as quantifiable peptides. We therefore focused below on isoforms with unique peptides for simplicity, and then inspected the non-unique results on a case-by-case basis. A number of proteins with multiple quantifiable isoforms show similar enrichment across isoforms, such as the atrium biased expression of fibulin-1 (FBLN1) canonical isoform and isoform −4; and ventricle heat shock protein family B (small) member 7 (HSPB7) canonical form and isoform −2. Other proteins show divergent distributions of isoforms. We highlight three cases below that illustrate the value of the workflow to mine additional chamber-enriched proteoforms from existing data.

**Figure 1:**
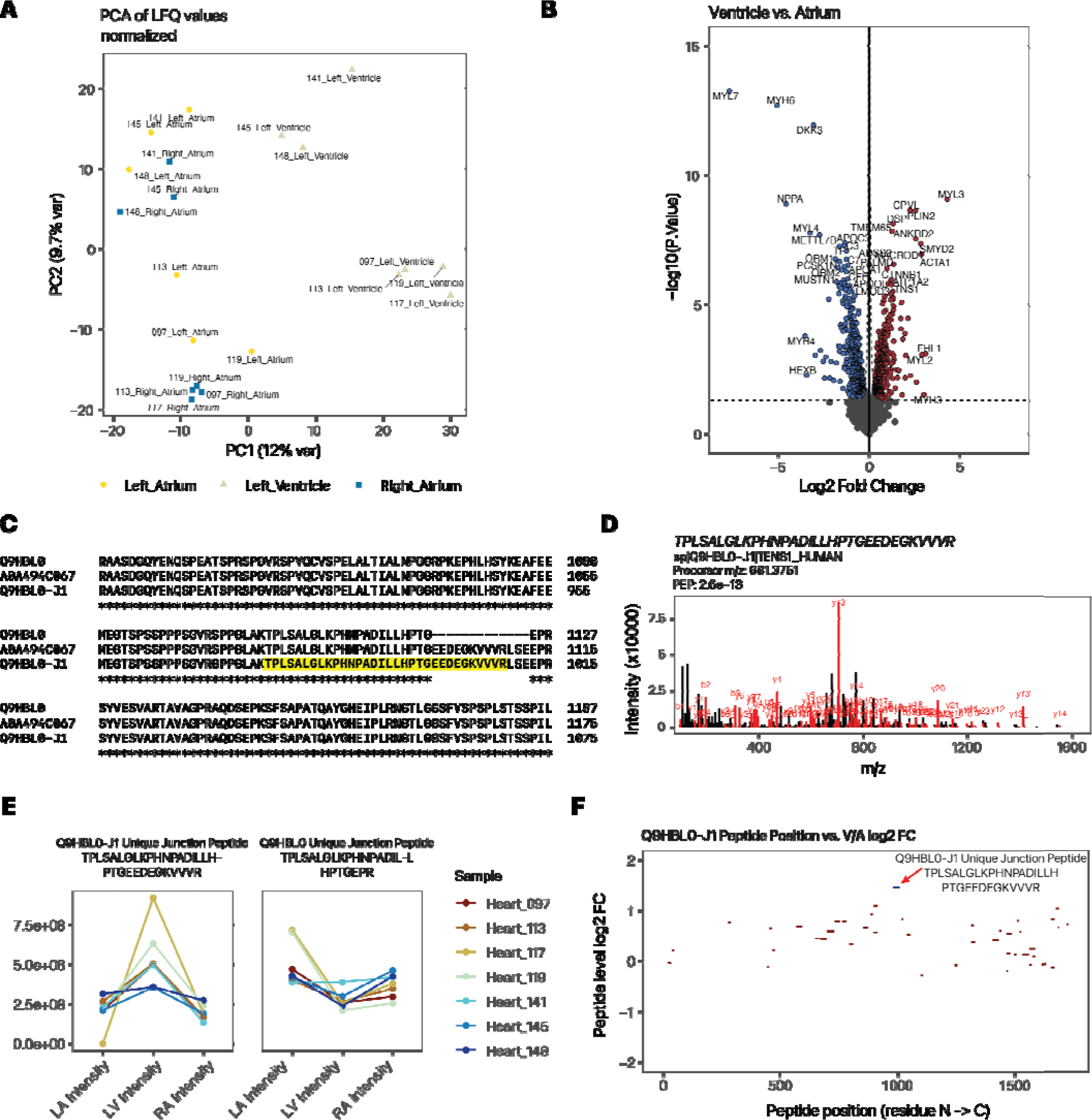
Quantification of atrium and ventricle distributions of protein isoforms show tissue usage preferences. **A.** PC1 vs. PC2 of custom database search, recapitulating the atrial-ventricular separation of total protein profiles. **B.** Volcano plot (log2FC vs. –log10 P) showing top differentially expressed proteins that are more highly expressed in the ventricle (right) or atrium (left) of the heart. **C.** ClustalOmega alignment of canonical tensin 1 (TNS1; Q9HBL0) and the custom translated isoform, which corresponds to the TrEMBL sequence A0A494C067 within the relevant splice junctions. **D.** Peptide spectrum match of the Tensin-1 isoform junction peptide. E. Line plots showing normalized protein expression in the left atrium (LA), left ventricle (LV), and right atrium (RA) of the splice junction-spanning peptide unique to the tensin-1 alternative isoform (left) vs. the corresponding junction-spanning peptide of the canonical form (right). Color: individual donor. **F.** Map of unique peptide by their residue location (x-axis) within the tensin-1 isoform sequence and their relative ventricle to atrium (V/A) expression ratio (y-axis). The isoform-specific junction peptide is in blue. Red peptides are shared with the canonical sequence. Only peptides with 1e8 or above LFQ intensity are included.

**Table 1:** List of proteins with two or more (canonical or alternative) isoforms quantified with unique peptides.

| Gene Name | Protein Name | UniProt | Log2 FC | B-H adj. P | Protein Pr. | Canonical Search log2 FC | Canonical Search adj. P |
| --- | --- | --- | --- | --- | --- | --- | --- |
| ADD3 | adducin 3 | Q9UEY8-2 | -0.72 | 4.12E-03 | 1.000 | -0.91 | 8.43E-04 |
| ATP2A2 | ATPase sarcoplasmic/endoplasmic reticulum Ca2+ transporting 2 | P16615-2 | -0.72 | 1.36E-03 | 0.999 | -0.99 | 2.14E-04 |
| CBFB | core-binding factor subunit beta | Q13951-2 | 0.54 | 3.16E-02 | 0.996 | 0.59 | 1.44E-02 |
| DGKZ | diacylglycerol kinase zeta | Q13574-1 | -0.82 | 3.16E-03 | 1.000 | -1.18 | 1.36E-04 |
| FBLN1 | fibulin 1 | P23142-4 | -1.37 | 5.97E-03 | 1.000 | -1.81 | 2.24E-03 |
| FBLN2 | fibulin 2 | P98095-2 | -0.92 | 4.02E-03 | 1.000 | -1.50 | 1.36E-04 |
| FHL1 | four and a half LIM domains 1 | Q13642-5 | 3.09 | 8.89E-03 | 1.000 | 1.40 | 3.76E-02 |
| HSPB7 | heat shock protein family B (small) member 7 | Q9UBY9-2 | 1.20 | 2.11E-03 | 1.000 | 1.32 | 2.22E-03 |
| IDH3B | isocitrate dehydrogenase (NAD(+)) 3 non-catalytic subunit beta | O43837-2 | -0.65 | 6.28E-02 | 1.000 | -0.38 | 1.27E-01 |
| LRRFIP1 | LRR binding FLII interacting protein 1 | Q32MZ4-4 | 0.69 | 2.02E-02 | 1.000 | 1.08 | 6.51E-02 |
| MRPS22 | mitochondrial ribosomal protein S22 | P82650-2 | 0.44 | 8.05E-02 | 0.990 | 0.02 | 9.25E-01 |
| MYBPC3 | myosin binding protein C3 | Q14896-J3 | 0.55 | 8.32E-03 | 1.000 | 0.46 | 1.34E-02 |
| NDUFV3 | NADH:ubiquinone oxidoreductase subunit V3 | P56181-2 | 0.34 | 3.47E-02 | 1.000 | 0.51 | 2.33E-01 |
| NSFL1C | NSFL1 cofactor | Q9UNZ2-5 | 0.50 | 1.53E-02 | 1.000 | 0.24 | 9.59E-02 |
| PDLIM3 | PDZ and LIM domain 3 | Q53GG5-2 | -0.55 | 7.03E-02 | 1.000 | -0.25 | 2.65E-01 |
| PDLIM5 | PDZ and LIM domain 5 | Q96HC4-3 | -1.23 | 1.25E-02 | 1.000 | 0.26 | 3.07E-01 |
| PDLIM5 | PDZ and LIM domain 5 | Q96HC4-6 | 0.79 | 2.11E-02 | 1.000 | 0.26 | 3.07E-01 |
| PFKFB2 | 6-phosphofructo-2-kinase/fructose-2,6-biphosphatase 2 | O60825-2 | 1.21 | 1.58E-02 | 1.000 | 1.76 | 1.36E-04 |
| PFN2 | profilin 2 | P35080-2 | -0.61 | 3.67E-03 | 1.000 | -0.93 | 1.51E-04 |
| PLEC | plectin | Q15149-8 | -0.34 | 7.26E-02 | 1.000 | 0.51 | 1.33E-02 |
| PNKD | PNKD metallo-beta-lactamase domain containing | Q8N490-2 | 0.69 | 8.41E-02 | 1.000 | 0.29 | 1.35E-01 |
| SLC25A3 | solute carrier family 25 member 3 | Q00325-2 | 0.63 | 8.30E-02 | 1.000 | 0.18 | 7.04E-01 |
| SORBS2 | sorbin and SH3 domain containing 2 | O94875-10 | 1.74 | 2.13E-03 | 1.000 | 1.09 | 4.12E-04 |
| SRL | sarcalumenin | Q86TD4-1 | 0.75 | 5.10E-02 | 1.000 | -0.11 | 6.41E-01 |
| STRN3 | striatin 3 | Q13033-2 | -0.50 | 2.56E-02 | 1.000 | -0.67 | 1.34E-02 |
| SYNPO | synaptopodin | Q8N3V7-2 | -0.62 | 9.56E-03 | 1.000 | 1.80 | 3.84E-03 |
| TNS1 | tensin 1 | Q9HBL0-J1 | 1.29 | 1.88E-04 | 1.000 | 0.43 | 2.02E-02 |
| TPM1 | tropomyosin 1 | P09493-10 | 0.41 | 7.58E-02 | 1.000 | 0.34 | 1.25E-01 |
| TTN | titin | Q8WZ42-6 | 0.53 | 1.08E-02 | 1.000 | 0.15 | 4.43E-01 |

#### PDLIM3

PDZ and LIM Domain 3 (PDLIM3) is a cardiac enriched protein localized to the Z disc. On UniProt SwissProt, PDLIM3 has three isoforms including the canonical −1 isoform and the −2 and −3 alternative variants. We highlight this case because in the original study, PDLIM3 was not reported to be significantly enriched in the atrium or the ventricle at the total protein level. In the re-analysis, we found PDLIM3 SwissProt reviewed isoform (Q53GG5-2) with a unique peptide GLIPSSPQNEPTASVPPESDVYR. The peptide was again identifiable from large data sets at PepQuery2. The isoform 2 was enriched in the atrium at 1.5 fold (adjusted p: 0.07). Noticeably, isoform 1 was enriched in the ventricle, although this difference is not significant (1.4-fold, adjusted p: 0.25). Nevertheless, the *isoform−1/−2 ratios* likely differ in the atrium and ventricle by ∼2.0 fold. The divergent ratios across the two isoforms likely explain its exclusion in the original study or the canonical search and attest to the caveats of protein inference.

#### Tensin-1

Tensin-1 is a protein that resides in the focal adhesion. A Tensin-1 isoform not present in SwissProt isoforms was identified from a sequence translated from RNA-seq splice junction data (Q9HBL0-J1). The in silico translated isoform differs from the canonical tensin-1 sequence on Uniprot (Q9HBL0-3) through an insertion of the amino acid sequence EEDEGKVVVRLSE between residues 1124 and 1125 of the canonical sequence. In addition, residues 1–125 at the N-terminus are missing due to the use of a different Ensembl annotated translation start site, but inference cannot be drawn on the full-length molecule due to the nature of short-read Illumina sequencing of the RNA-seq data, which does not allow the read variants at different regions of the transcripts to be confidently linked. The insertion is also found in the TrEMBL sequence A0A494C067, which is also missing the first 25 N terminal residues but is not documented within SwissProt canonical + isoform sequences (**Figure 1C**).

In the experiment, the Q9HBL0-J1 isoform was identified by two partially overlapping unique peptides (TPLSALGLKPHNPADILLHPTGEEDEGK and TPLSALGLKPHNPADILLHPTGEEDEGKVVVR), which span the splice junction containing the unique insertion (**Figure 1D spectrum**). The -J1 isoform was significantly enriched in the ventricle (V/A ratio: 2.4-fold, adjusted P 1.8e–4). In the canonical database search, Tensin-1 is significantly enriched in the ventricle but in a more modest manner (V/A ratio: 1.4-fold, adjusted p: 0.02) and similarly in the Lindscheid et al. report (V/A ratio: 1.4-fold, p: 0.001) which suggests the reported fold change at the gene level may be due to a combined contribution of canonical and variant proteins. Closer inspection of the quantitative data of the corresponding splice junction peptides show that the isoform-specific junction peptide exhibits a strong ventricular enrichment in expression that is not present in the corresponding canonical form (**Figure 1E**). Moreover, the proteoform-specific peptide has a V/A ratio that is placed ∼2.95 standard deviations from the mean of peptides that are shared with canonical tensin-1 (**Figure 1F**). Therefore, we surmise that a tensin 1 isoform can be found at both the peptide level and protein level and is significantly enriched in the ventricle.

The splice variant peptides span the junction of the variant region of the isoforms (residues 878 to 1009) and do not match to any sequence within SwissProt when allowing up to two residue mismatches. As mentioned above, the peptide sequences are also contained within TrEMBL sequence A0A494C067 (residues 1078 to 1109) but are otherwise uncharacterized to our knowledge. We used two complementary approaches to validate the identification. First, we compared the custom protein database search with a different search engine and filtering pipeline (Comet (29) and Percolator (30)). We found the two sequences to be confidently identified at Percolator with multiple-testing-adjusted posterior error probability < 0.01 in 57 peptide-spectrum matches (PSMs) across 20 samples, with 56 of these coming from the longer sequence. None of the spectrum matches could be identified confidently in a corresponding search using a canonical database (posterior error probability of the best-fit target peptides: 0.89 – 0.99), suggesting the isoform-containing custom database provides a substantial increase in explanatory power of the mass spectra. Secondly, we queried millions of additional public mass spectrometry data on non-cancer tissues using a targeted peptide search engine PepQuery2 (46), which allows novel peptide sequences to be identified across over a billion pre-indexed mass spectra. The longer proteoform-unique peptide (TPLS…VVVR) is again confidently identified from a spectrum (01296_E02_P013201_S00_N13_R1:29599:5) in the “29_healthy_human_tissues” (PXD010154) dataset (hyperscore 83.29) and moreover explains the spectrum considerably better than the best PSM from the reference database search (hyperscore 18.97) or unrestricted modification-based searching (hyperscore 28.95) (**Supplemental Figure S1A**). Likewise, the shorter peptide specific to the proteoform (TPLS…DEGK) is confidently identified in a spectrum (Instrument1_sample13_S1R9_072116_Fr12:38150:5) in the “GTEx_32_Tissues_Proteome” dataset (PXD016999) (hyperscore 99.63) and again outperformed the reference database search (hyperscore 54.28) and the unrestricted modification-based search (hyperscore 72.34) of the same spectrum (**Supplemental Figure S1B**). Taken together, the results support bona fide identification of the non-canonical peptides.

#### PDLIM5

Other isoforms were observed that present less clear-cut examples of atrial or ventricular enrichment, buthighlight the intrinsic complexities in mapping isoform peptides through bottom-up proteomics. One example is PDZ and Lim Domain 5 (PDLIM5). The *PDLIM5* gene has a complex splicing pattern, and splicing errors have been linked with left ventricle non-compaction. With the complexity of its 7 overlapping SwissProt isoforms, no unique peptides were quantified in the re-analysis. However, in the unique+razor peptide quantification, we found the isoforms −3 and −6 to be identifiable by unique peptides with protein probability > 0.99; whereas the -J1 and canonical −1 isoforms were both identified with protein probability > 0.95 but were indistinguishable from their component peptides. However, as it is known that in most proteins in most tissues, the canonical forms are expressed at a higher level than alternative isoforms, hence we interpreted the peptides to belong to the canonical form. The canonical form showed no significant V/A enrichment (1.4 fold, adjusted p: 0.13); whereas isoform −6 is modestly enriched to the ventricle (1.6 fold, adjusted p: 0.025) but isoform −3 is enriched to the atrium (2.2 fold, adjusted p: 0.012).

#### Validation data

To corroborate the quantitative findings on tensin-1 and PDLIM-3, we performed MSFragger/IonQuant label-free quantification on a separate deep mass spectrometry data with atrial and ventricle samples (PXD006675 Doll et al.; LA, RA, LV, n=3 male hearts) **(Supplemental Table S3 Doll limma results)**. The results corroborate the atrium enrichment of the PDLIM3-2 isoform (Q53GG5-2; 2.5 fold, adjusted p: 2.0e–3), the ventricular enrichment of the tensin -J1 isoform (Q9H8J0-J1; 1.6 fold, adjusted p: 0.08) and the ventricular enrichment of the PDLIM5-6 isoform (Q96HC4-6; 1.5 fold, adjusted p: 0.08), which lends support to their tissue-biased usage across two independent cohorts.

### Inferring the molecular functions of chamber-biased splice variants

We next considered the potential molecular function of the chamber-biased proteoforms. First, we explored the potential function of the PDLIM3-2 isoform (ALP-H). Canonical PDLIM3 is a Z disc protein that contains an N-terminal PDZ domain (residues 8 – 84), followed by an extended unstructured region (residue 111 to 238) that include a Zasp-like motif (residue 184 – 209), and followed by a structured LIM domain near the C-terminus (residues 292 – 351). The PDZ domain is known to interact with α-actinin 2 (ACTN2). The AlphaFold2 proposed structures for PDLIM3 and PDLIM3-2 showed overall pLDDT of 65.2 and 68.3. By comparison, the PDLIM3-3 isoform which omits the C-terminal LIM domain has an overall pLDDT of 66.9. As expected, the N-terminal and C-terminal domains in PDLIM3 and PDLIM3-2 are associated with high confidence structural prediction (pLDDT > 90) whereas the internal regions with low confidence of structure prediction (pLDDT < 60) (**Figure 2A**). The predicted aligned errors (PAE) graph of PDLIM3 protein showed that sequence position S148–E216 had lower confidence in the structure folding, which intersects the variant sequence portion from the canonical isoform.

**Figure 2:**
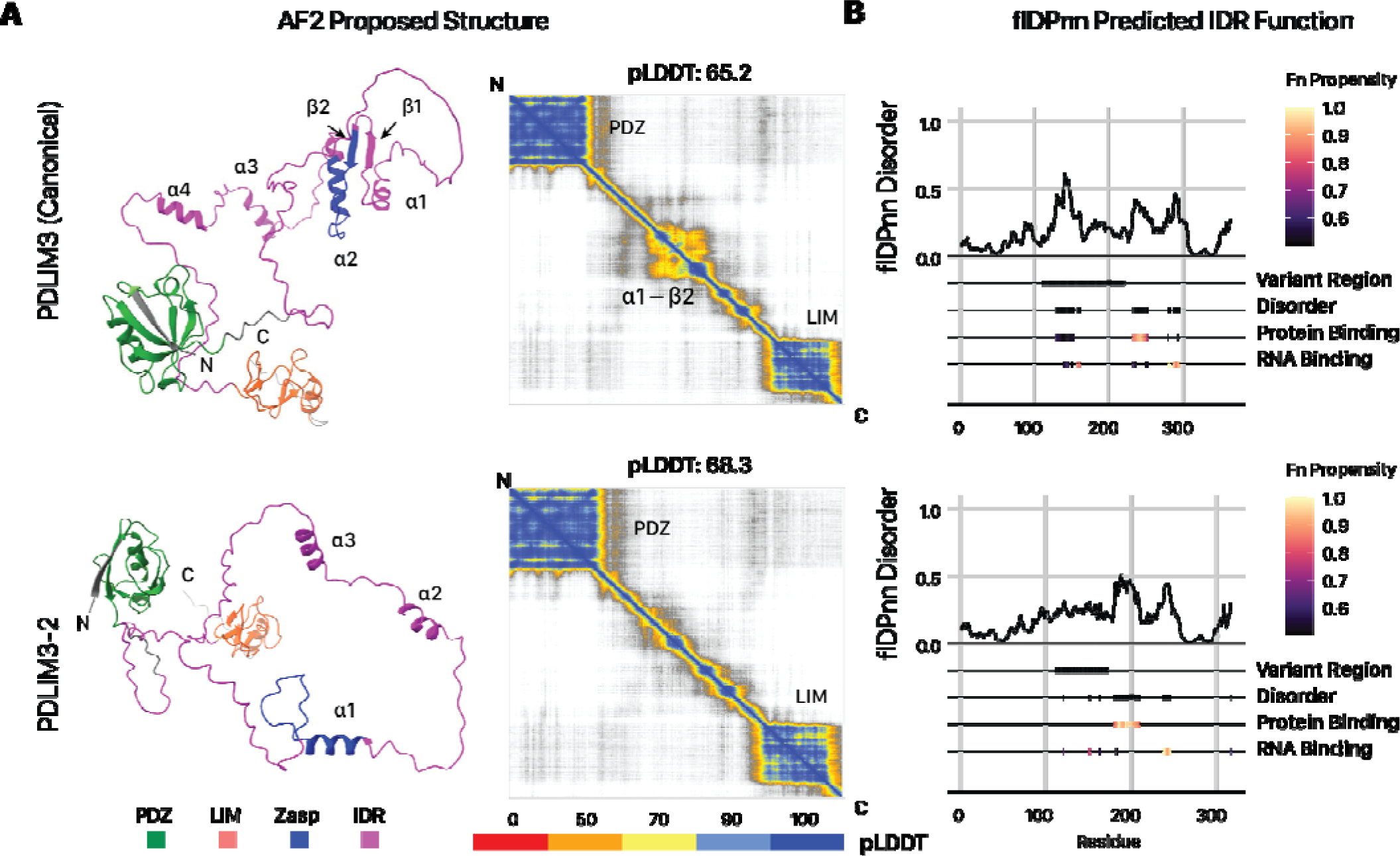
The PDLIM3-2 isoform is associated with altered structure and IDR predictions. **A.** AlphaFold2 proposed structure for PDLIM3 canonical (top) and the PDLIM3-2 alternative (below) isoforms and the prediction alignment error (PAE) matrix (right)**. B.** flDPnn predicted sequence disorders and functional features. Darker colors represent higher propensity for each function. The PDLIM3-2 isoform shows greater protein binding propensity within the internal IDR that overlaps with the isoform-variant region.

PDLIM3-2 differs from PDLIM3 through a variant region (111–224) that intersects with the extended internal IDR between the PDZ and LIM domains. In the IDR, the PDLIM3 sequence has α1, α2 and β1 & β2 folds that are missing in the isoforms due to the deletion and variation in the sequence (**Figure 2B**). Moreover, the variant regions intersect with an InterPro unknown function domain DUF4749 with small ZASP motif sequence (K184 to Q209 of canonical). ZASP motifs are known to function in the efficiency of PDZ binding to α-actinin for increasing the integrity in the Z-line (47, 48). As the ZASP-like motif sequences between canonical (PDLIM3) and non-canonical (PDLIM3-2) isoforms differ from each other, we asked whether the variant IDR may exert an effect on PDLIM3 protein-protein interaction. Consistent with this, flDPnn prediction for the isoforms of PDLIM3 confirmed that the variant regions show strong changes in disorder propensity and classification, with an extended region that is strongly predicted for protein binding (**Figure 2C**). To corroborate this predicted function in the protein interaction, we further subjected the AlphaFold2-proposed structure of each isoform to analysis by ScanNet, a deep learning model for structure-based protein interaction predictions, which showed that PDLIM3-2 has on average higher protein binding propensity than the canonical (**Figure 3A**). Finally, AlphaFold-Multimer prediction predicts a clear increase in the interaction between alpha actinin (ACTN2) with PDLIM3-2 (**Figure 3B**). Taken together, multiple predictions algorithms corroborate to suggest a role of PDLIM3-2 in altering biological function in the variant IDR including potentially difference in protein-protein interactions including potential strengthening of actinin interactions, which may be tested in future experiments.

**Figure 3:**
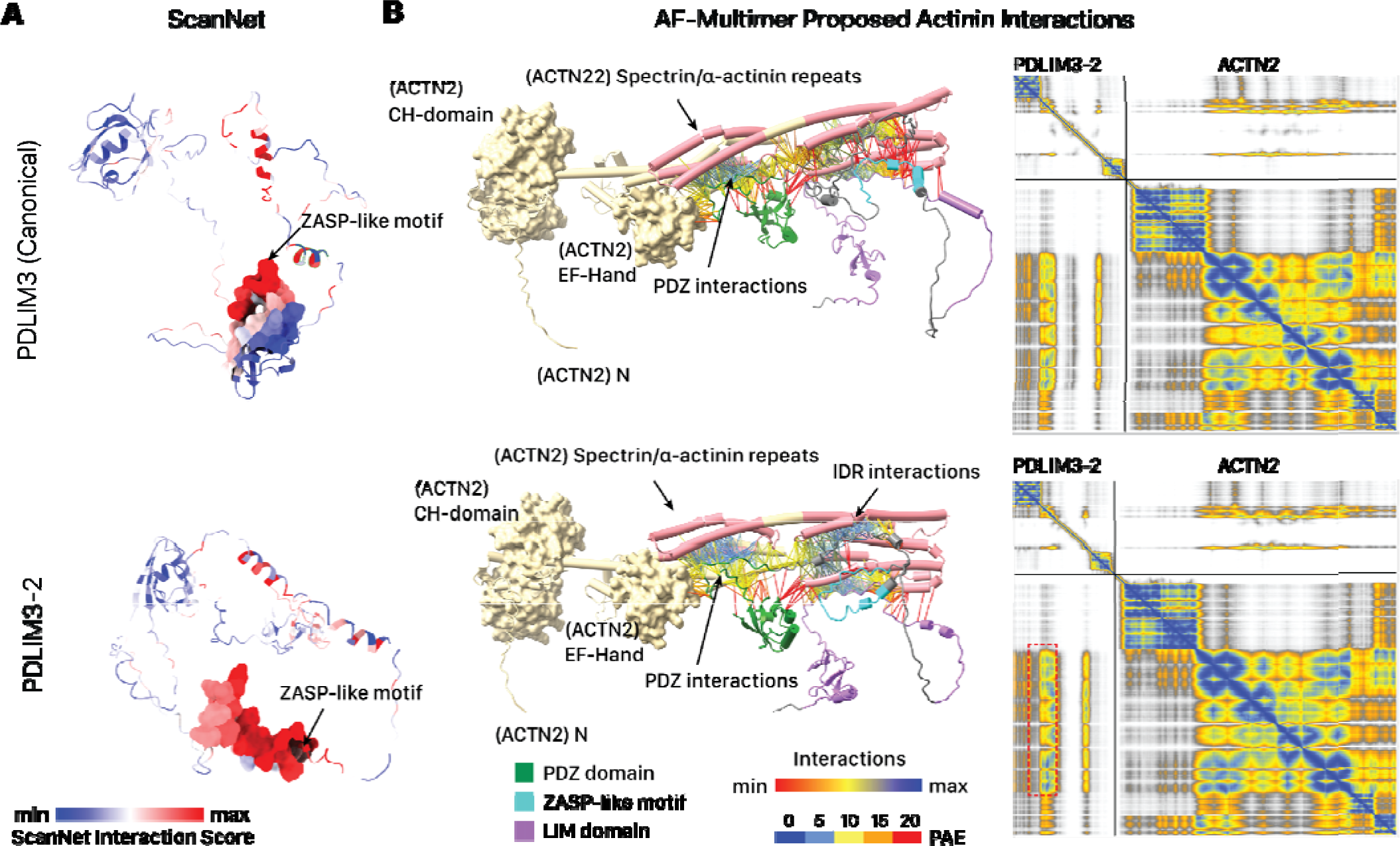
The PDLIM3-2 isoform is associated with differences in protein-protein interaction predictions. **A.** ScanNet prediction of protein-protein interaction surfaces show higher interaction propensity in the PDLIM3-2 isoform than the canonical sequence. Red: higher ScanNet interaction score. **B.** AlphaFold multimer prediction of PDLIM3 (top) and PDLIM3-2 (bottom) binding with alpha actinin (ACTN2). The upper structure is human ACTN2 and lower structure is PDLIM3. Interaction linkages and PAE matrix colors represent PAE for residue pairs at interfaces displayed by ChimerX; blue linkages represent higher interaction scores. Protein-protein interaction links show PDLIM3 binding with the spectrin/alpha-actinin repeats (cylinder), with increased binding in PDLIM3-2 than canonical PDLIM3 and due to sequence variation in isoform (red dashed box on the PAE graph).

The isoforms of PDLIM5 share the PDZ domains, whereas the PDLIM5-3 isoform omits the LIM domain and PDLIM-6 has greater protein propensity in the IDR region (**Supplemental Figure S3**). In contrast, few conclusions can be currently drawn about the function of the tensin-1 variant. The variant splice junction intersects with the canonical protein residues 1124–1135, which are not adjacent to any annotated domains or sequence features on UniProt. Structural prediction by AlphaFold2 fails to return a stably folded structure in the majority of either the canonical or variant proteins, aside from the tensin-type, SH2, and PTB domains. Consistent with this, using the disorder prediction algorithms IUPred3 (49) and Metapredict v2 (42) we find that most regions including from residues 439 to 1552 are in an extended unstructured region. Given the function of tensin-1 as a scaffold protein for adhesion signaling (50), it is likely that the unstructured regions participate in protein interactions. The potential functions of the PDLIM5 and tensin-1 splice variants therefore await further investigation.

### Evidence for intrinsically disordered region remodeling among alternative isoforms

We then investigated whether there are proteome wide changes in IDR functions among all expressed alternative isoforms in the heart. Therefore we considered both data sets (PXD006675 and PXD008722) including mass spectra from the LA, LV, and RV of 7 human donors in Linschied et al. and the LA, LV, RA, and RV of 3 donors in Doll et al. We prioritize a set of high-confidence non-canonical isoforms, hich are supported by unique peptides and are identified at >99% Philosopher Protein Probability as well as Posterior Error Probability ≤ 0.01 in Comet/Percolator (**Figure 4A**). This shortlisted 216 high-confidence non-canonical protein isoforms (**Supplemental Table S4**), including 164 SwissProt isoforms, and 52 isoforms that are not documented in SwissProt but are included by virtue of the custom RNA-seq translated database.

**Figure 4:**
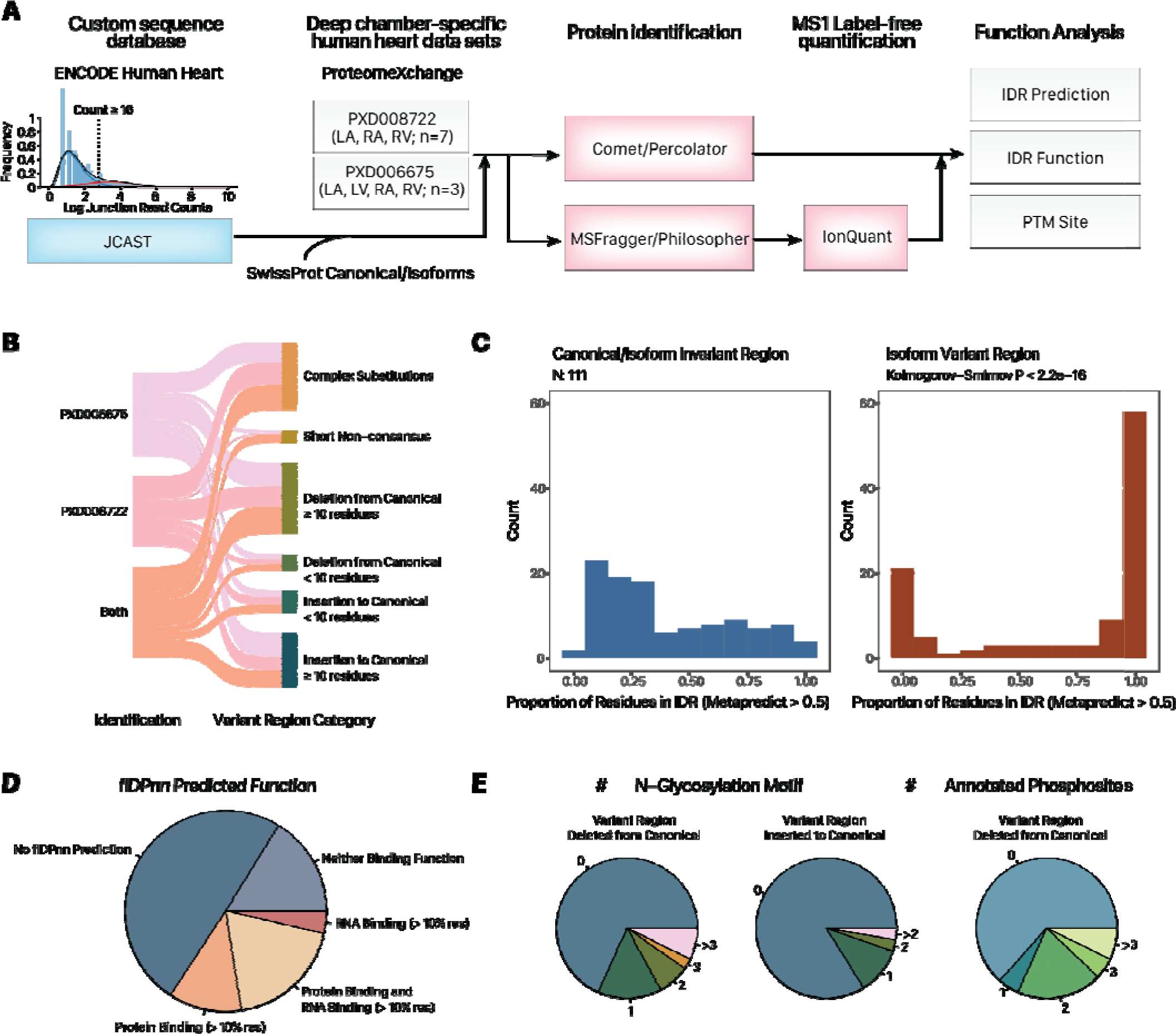
Translated alternative splicing protein isoforms intersect with intrinsically disordered regions with predicted function. **A.** Proteogenomics workflow. Inset histogram shows the distribution of splice junction read counts in the ENCODE RNA sequencing data. Only splice junction pairs with 16 or more total reads are translated to proteins in silico. **B.** Sankey diagram showing the number of identified protein isoforms with unique peptides. 111 of the isoforms containing a simple insertion or deletion of 10 or more residues from the canonical sequences are prioritized for analysis. **C.** Proportion of residues within canonical/alternative invariant and variant regions that are annotated to be within IDRs (MetaPredict v2 score > 0.5). Kolmogorov-Smirnov P < 2.2e–16. **D.** Pie chart showing the percentage of IDRs within isoform variant regions that are annotated to be protein binding and/or RNA binding. **E.** Pie charts showing the number of non-canonical isoforms with variant regions intersecting with various numbers of N-glycosylation sequence motifs and annotated phosphorylation sites.

To gain insight into the potential function of the identified isoforms, we prioritized 111 isoforms that lend themselves to interpretation because their variant sequences differ from their canonical counterparts by a simple variant region comprising a single insertion/deletion of a >10 aa, as recognized by a local alignment algorithm (**Figure 4B**). We then used MetaPredict which uses multiple algorithms to predict and vote on whether any amino acid belongs to an IDR region in a protein sequence. Among the 111 examined isoforms, we found that over two-third had variant regions that contained 50% or more residues predicted to be disordered, which is significantly higher than the less than one third in the invariant regions (i.e., sequences not altered by the alternative isoform) (Komogorov Smirnov p < 2.2e–16) (**Figure 4C; Supplemental Table S5**). Our findings show that the translated protein isoforms are significantly enriched in IDRs within regions that differ from their canonical counterparts.

We next used the deep learning model flDPnn as above to predict the function of the disordered sequences in all variant regions (inserted or deleted from canonical sequences). flDPnn has a more stringent requirement for disordered regions than Metapredict and only predicts function within assigned IDRs. Among the 56 isoforms with any flDPnn functional predictions, over two-thirds or 38 contain residues that are confidently assigned to have RNA-binding or protein binding function whereas over one-third or 21 are associated with both RNA-binding and protein-binding functions (**Figure 4D**). Across all 111 alternative isoforms with simple variant regions (i.e., single insertion or deletion of ≥ 10 residues from the canonical sequence), the predicted IDR sequences within variant regions are significantly more likely to contain flDPnn-predicted protein binding (Kolmogorov-Smirnov p: 4.5e–9) and RNA binding function (p: 3.3e–10) than predicted IDR sequences across canonical-alternative shared regions (**Supplemental Figure S4**). The analysis reveals that translated protein isoforms not only impact IDR distributions of cardiac proteins substantially, but are moreover significantly enriched in IDR regions associated with predicted function. In parallel, the translated alternative isoforms show substantial overlaps with PTMs, with over one-third of isoforms associated with one or more annotated phosphorylation sites at the removed region of the canonical sequence (**Figure 4E**).

Lastly, the below examples illustrate the extent to which alternative splicing derived variant regions can intersect with protein features of interest. Isoform 2 of NADH:ubiquinone oxidoreductase subunit V3 (NDUFV3) includes a long insertion at residue 57–417 with predicted IDR and protein binding propensity, an N-glycosylation motif, and multiple annotated phosphorylation sites (**Figure 5A**). For sarcalumenin (SRL), the −2 isoform has been annotated as the canonical by Uniprot; the −1 alternative isoform contains a long inserted IDR region with protein binding propensity and an N-glycosylation motif (**Figure 5B**). Finally, for microtubule-associated protein 4 (MAP4); an RNA sequencing derived isoform J6 with long inserted IDR shows multiple N-glycosylation motifs (**Figure 5C**). Similar observations were made for isoforms that contain removed regions from the canonical sequences, overlapping with IDRs and phospho-IDRs (**Supplemental Figure S5**). Altogether, these examples strongly support the notion that alternative splicing can produce translated stable protein products with considerable potential for modified function through PTM and IDR remodeling.

**Figure 5:**
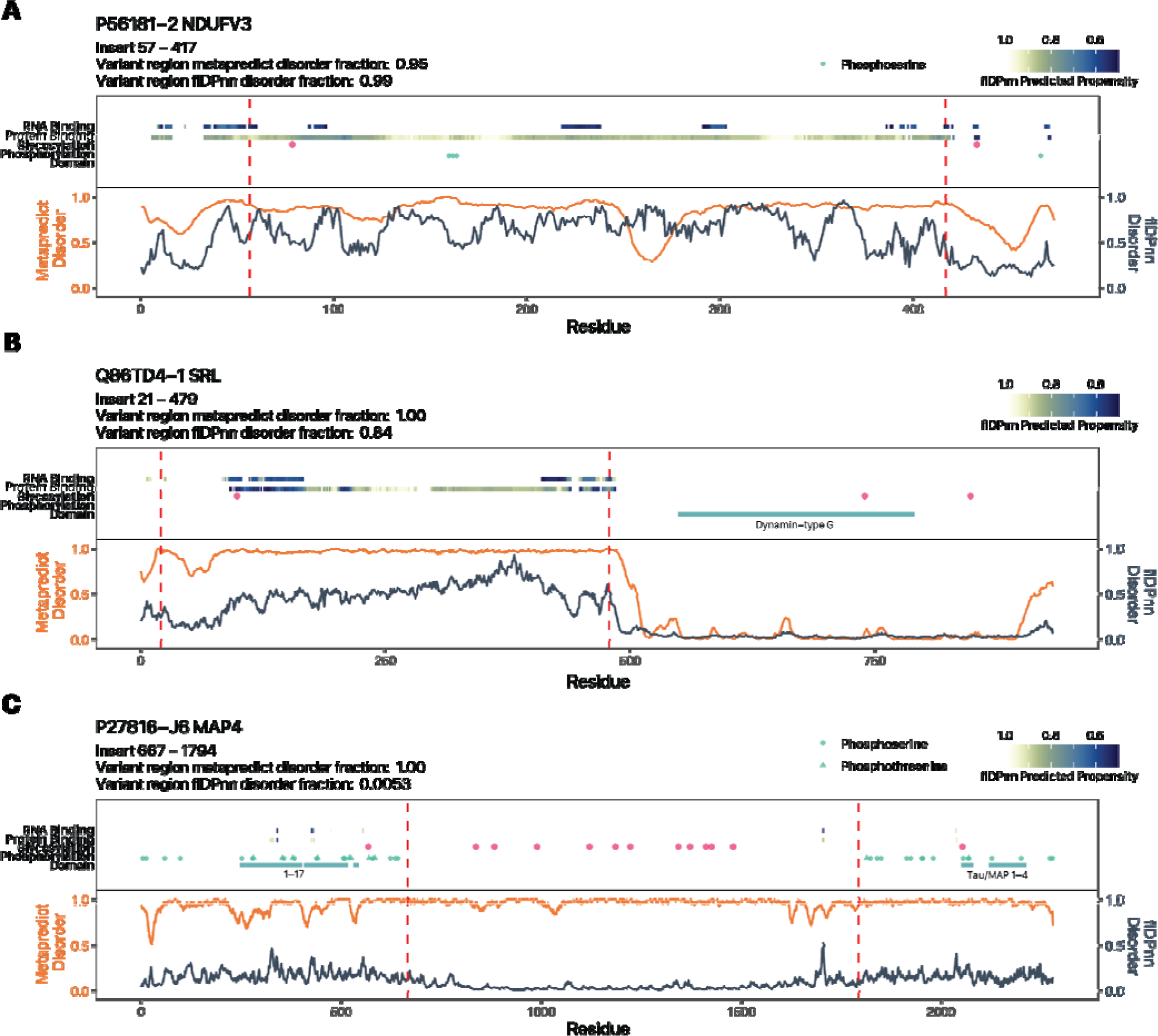
Alternative isoforms with inserted variant regions show evidence of IDR remodeling. Three isoforms with inserted variant regions over the canonical sequences are shown: **A.** NDUFV3-2; **B.** SRL-1; **C.** MAP4-J1. Orange trace represents MetaPredict v2 disorder score. Blue trace represents flDPnn disorder score. Red dashed lines demarcate the variant regions between the canonical and alternative isoforms. Tracks from bottom to top: UniProt/InterPro annotated domains; UniProt annotated phosphorylation sites; NXS/T N-glycosylation motifs; flDPnn protein binding propensity; flDPnn RNA binding propensity.

## Discussion

The function of splice variant proteins remains poorly understood and their overall importance to the diversity of proteoforms has been disputed in the literature (5). Computational advances have allowed more proteoforms to be conclusively identified, and the heart has emerged as one of the organs that show more proteoform diversity due to splicing (9). Recently, structural prediction methods such as AlphaFold2 have been used to determine the sequence-structural-function relationships of protein isoform sequence (51), however the focus of such work has continued to be finding well-folded structure, e.g., transcript isoforms with higher overall pLDDT than the annotated canonical forms despite instances of known functional isoforms failing to fold into stable structures using current state of the art methods (51). This is consistent with known overlap between alternative isoforms with unstructured regions outside of stable domains at the transcript levels (5, 52), and suggests new methods are needed to find sequence-structure-function relationships. Prior work using exogenously expressed yeast two-hybrid or co-immunoprecipitation experiments (2, 53) and domain-based computational annotations (54) have shown that alternative isoforms may function to rewire protein-protein interaction capacity. Here we provide evidence applying new functional prediction tools that among hundreds of endogenously translated stable protein products of alternative splicing transcripts, altered protein-protein interaction capacity likely arises in part through the remodeling of IDR functions. Translated protein isoforms that differ from their canonical counterparts via the insertion or deletion of variant regions (e.g., through cassette exons) are highly enriched in IDRs, which are broadly associated with predicted function in protein binding, and at least in one examined case (PDLIM3-2), show different co-folding with a known interactor (ACTN).

Knowledge on proteome makeup differences across chambers can further current understanding of chamber specific diseases. It may also find utility in iPSC applications such as to act as molecular markers for protocols that focus on making atrial or ventricular cardiomyocytes. A limitation of the study however is that the samples from PXD008722 were collected from male patients with mitral valve regurgitation and dilated left atrium but no history of atrial fibrillation. Hence, the observed tissue usage differences may not generalize to female hearts, and moreover left atrial pathology may present a confounding factor for proteoform expression. However, as noted in the original publication by Lindscheid et al., over 98% of the atrial proteomes were not different between the left atrium and the undilated right atrium, and combining left and right atrial samples allowed an effective atrial vs. ventricular comparison. We therefore took the same approach of combining left and right atria to generate hypotheses on atrial vs. ventricular protein expression. The results of this study support the hypothesis that additional atrial and ventricular differences may be found through in-depth protein analysis. These signatures may come in several forms: novel peptides/proteins undocumented in commonly used protein databases, and span genes including PDLIM3 and PDLIM5 that have been associated with cardiomyopathy (55) thus highlighting the value of resolving protein isoforms and considering the nuances in protein inference in proteomics experiments.

In summary, this study (1) demonstrates a computational proteogenomics workflow to re-quantify mass spectrometry data can uncover additional proteoform information, (2) presents evidence of tissue usage preference for tensin-1 and PDLIM3/5 isoforms across the human heart atrium and ventricle, and (3) employs state-of-the-art prediction algorithms to explore the potential functional impact of alternative splicing through IDR and PTM remodeling. With the accelerating growth of the volume and depth of proteomics data being made available in recent years, we expect the overall approach outlined here to be useful for finding new regulatory principles of proteomes.

## Author Contributions

BP, SB, EL performed original investigations. BP, EL, MPL drafted the manuscript. BP, SB, AB, EL, DN, MPL revised the manuscript. EL and MPL handled funding.

## Acknowledgments

This work was funded in part by NIH award R35-GM146815 and the University of Colorado School of Medicine Translational Research Scholars Program to E.L.; and NIH awards R21-HL150456, R01-GM144456, and R01-HL141278 funding to M.P.L.

## Supplemental Tables and Figures

**Supplemental Table S1:** limma results from canonical database search using MSFragger/Philosopher in PXD008722

**Supplemental Table S2:** limma results from JCAST canonical + isoform database search using MSFragger/Philosopher in PXD008722

**Supplemental Table S3:** limma results for alternative isoforms quantified in validation data

**Supplemental Table S4:** All identified isoforms by Comet/Percolator and MSFragger/Philosopher

**Supplemental Table S5:** Annotated protein features in identified isoforms.

**Supplemental Figure S1:**
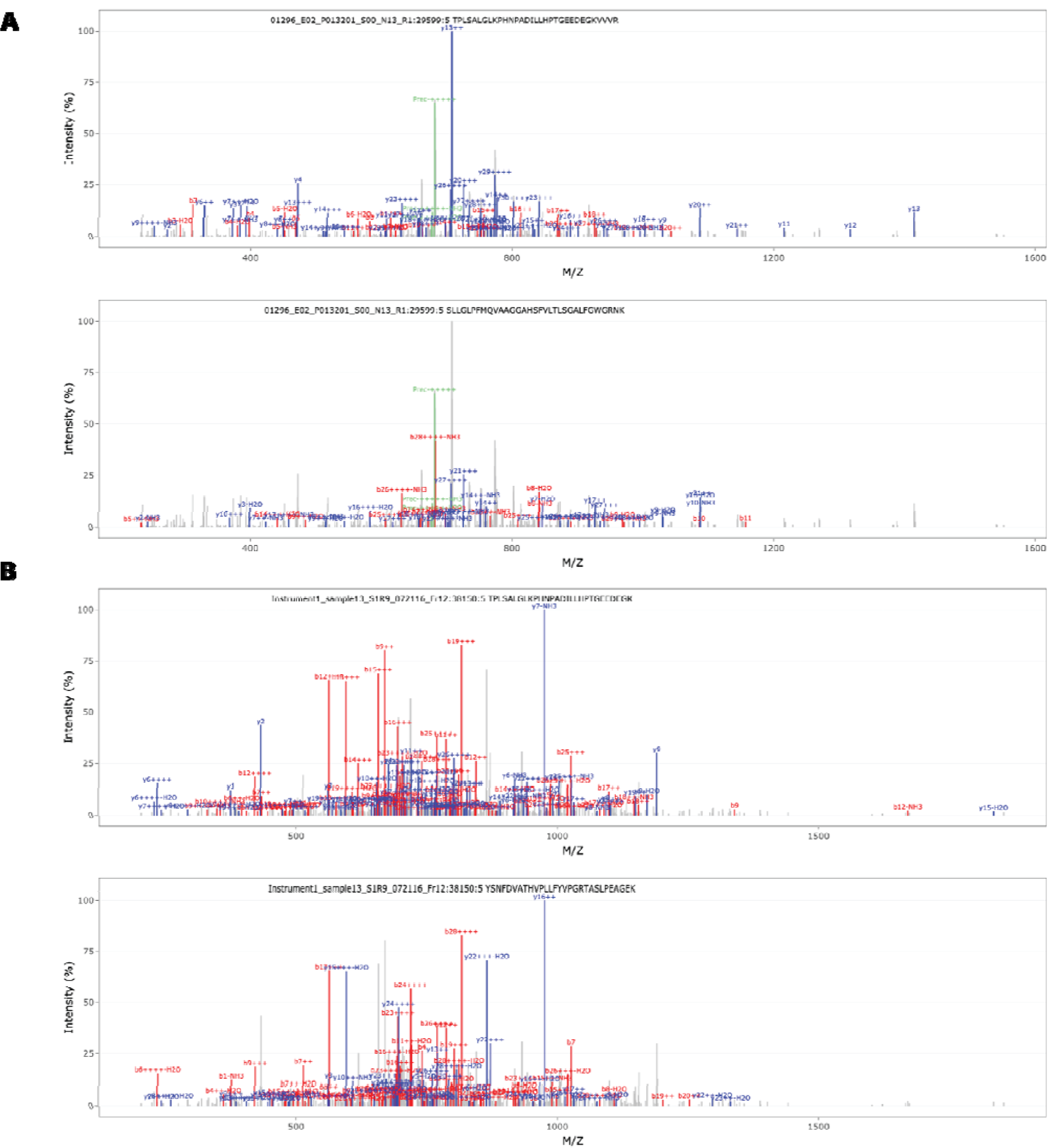
PepQuery2 peptide-spectrum matches for two undocumented tensin-1 -J1 isoform peptides. **A.** Peptide spectrum match of the splice junction peptide TPLSALGLKPHNPADILLHPTGEEDEGKVVVR against the “29_healthy_human_tissues” (PXD010154) dataset (hyperscore 83.29) (top) and the corresponding peptide spectrum match from the canonical database search (bottom). **B.** Peptide spectrum match of the splice junction peptide TPLSALGLKPHNPADILLHPTGEEDEGK against the “GTEx_32_Tissues_Proteome” (PXD016999) dataset (hyperscore 99.63) (top) and the corresponding peptide spectrum match from the canonical database search (bottom).

**Supplemental Figure S2:**
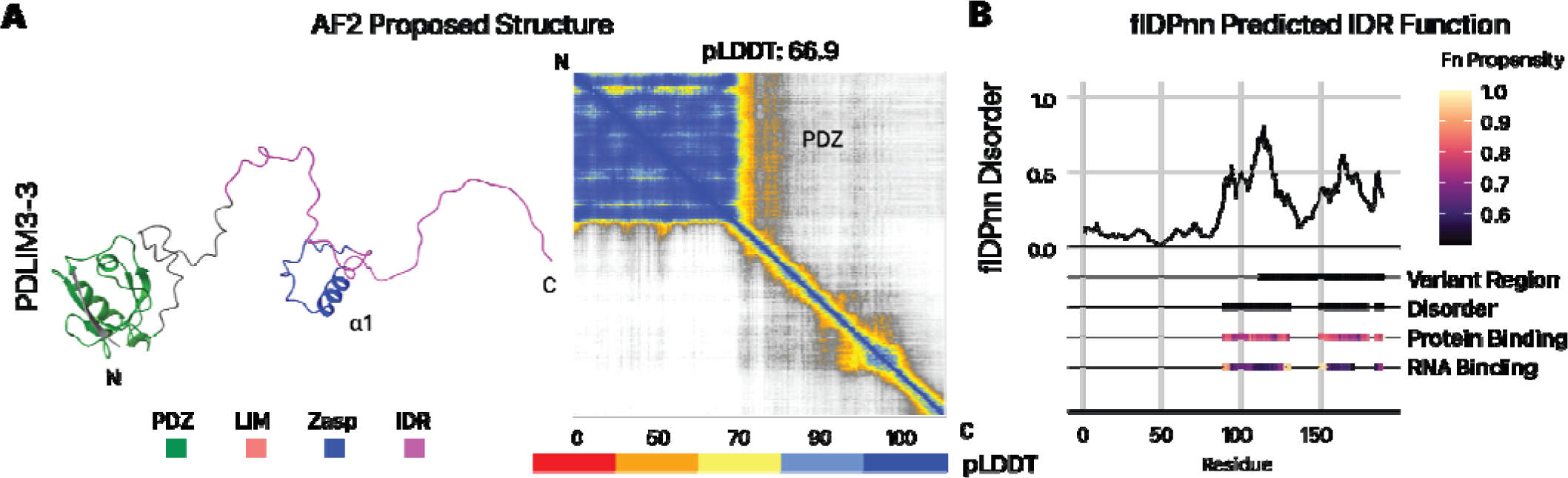
**A.** AlphaFold2 proposed structure and **B.** flDPnn predicted sequence disorders and functional features for the PDLIM3-3 isoform. See legends for Figure 2 for additional details.

**Supplemental Figure S3:**
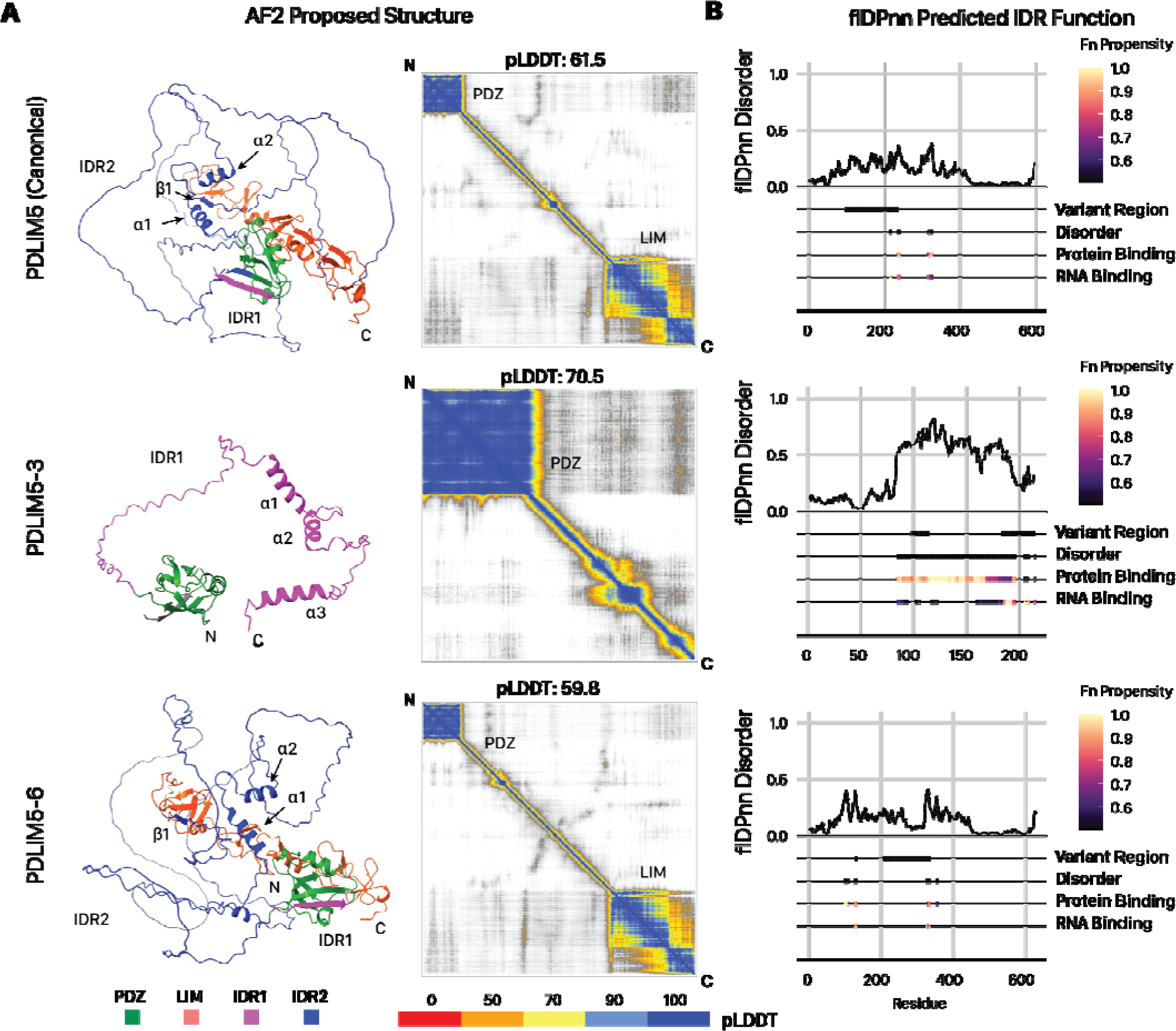
**A.** AlphaFold2 proposed structure and **B.** flDPnn predicted sequence disorders and functional features for the PDLIM5 sequence and the −3 and −6 alternative isoforms. See legends for Figure 2 for additional details.

**Supplemental Figure S4:**
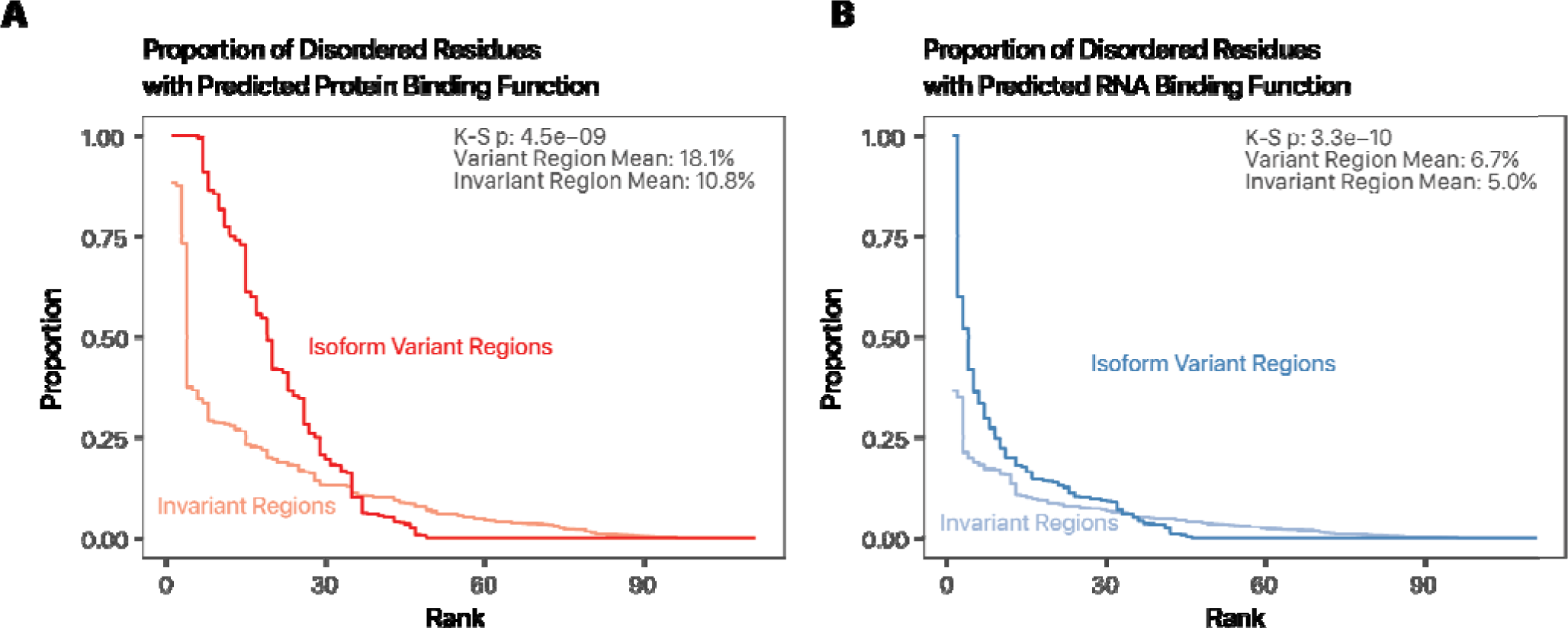
Proportion of residues within flDPnn predicted IDRs that are associated with **A.** protein binding or **B.** RNA binding function. K-S: two-sample Kolmogorov-Smirnov test. Darker line: IDR residues within isoform variant regions; lighter line: outside the isoform variant regions.

**Supplemental Figure S5:**
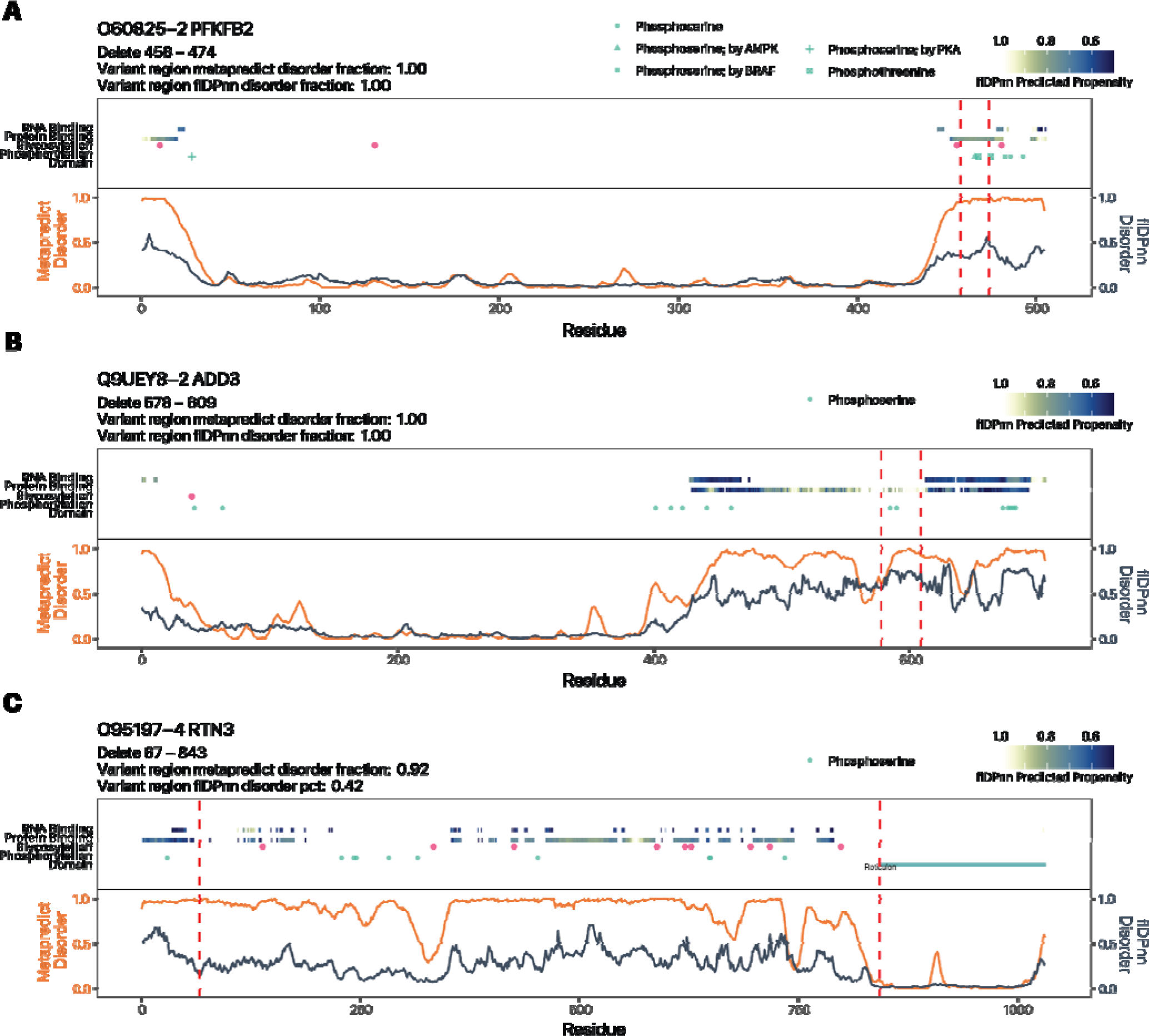
Alternative isoforms with deleted variant regions from canonical sequences show evidence of IDR remodeling. Three isoforms with variant regions removed from the canonical sequences are shown: **A.** PFKFB2-2; **B.** ADD3-2; **C.** RTN3-4. See legends for Figure 5 for additional details.

## Notes

### Competing Interest Statement

The authors have declared no competing interest.

